# Extreme conservation of miRNA complements in Opisthorchiids

**DOI:** 10.1101/146654

**Authors:** Vladimir Y. Ovchinnikov, Viatcheslav A. Mordvinov, Bastian Fromm

**Affiliations:** Department of Human and Animal Genetics, The Federal Research Center Institute of Cytology and Genetics The Siberian Branch of the Russian Academy of Sciences, Prospekt Lavrentyeva 10, Novosibirsk, 630090, Russian Federation; Department of Tumor Biology, Institute for Cancer Research, The Norwegian Radium Hospital, Oslo University Hospital, PO Box 4950, Nydalen, N-0424 Oslo, Norway

**Keywords:** *Opistorchis*, *Clonorchis*, microRNAs, biomarkers, host-parasite interaction

## Abstract

MicroRNAs (miRNAs) are key players in parasite-host communication and potential biomarkers for the detection of parasitic infections from host blood. Consequently, it is crucial to precisely know the miRNA complements of medically important agents such as the liver flukes of the Opisthorchiidae. Using publicly available and new datasets we curated and reannotated the surprisingly small and variable miRNA complements previously described for *Opistorchis viverrini, O. felineus* and *Clonorchis sinesis.* We find three highly similar miRNA complements with 53 identical and two miRNA genes with species specific sequences that signify a set of potential biomarkers and promising candidates for further investigations.

---

Liver flukes are parasitic flatworms that affect many different species of economic relevance and also infect humans. The medically most important group of liver-flukes is the Opisthorchiidae with Eurasian wide distributed species *Opisthorchis felineus, Opisthorchis viverrini* and *Clonorchis sinensis* (Figure 1) [1]. They are agents of human infections with Clonorchiasis / Opisthorchiasis transmitted by raw or undercooked fish and it is estimated that at least 1.2 million people worldwide are infected with *O. felineus,* 10 million with *O. viverrini* and 35 million with *C. sinensis* [2]. In humans, the infections are characterized by long durations, can occur with frequent exacerbations or without symptoms, and they may contribute to liver cancer development [3, 4]. Because Opistorchiids are classified as biocarcinogens, they came into the focus of various “OMICS”-studies that aimed at their characterization and, ultimately, identification of biomarkers towards the development of treatments [5, 6]. One important class of potential biomarkers is microRNAs (miRNAs) and we and others have recently shown their importance in host-parasite communication and immune-modulation [7,8,9]. miRNAs are small non-coding RNAs that are post-transcriptional gene regulators with important roles in many biological processes [10]. Previously, miRNA complements of the 3 opisthorchiids were published and they showed a surprisingly variable number of miRNA genes (between 16 and 18 conserved and between 20 and 43 novel genes) that was unexpected [11, 12]. The number of identified genes was also very low given the predictions from studies on miRNA evolution in flatworms [13] but most importantly no abundantly expressed miRNA was identified that showed sequential differences in all 3 species. The goal of our study was therefore first to curate and reannotate the miRNA complements of *O. viverrini, O. felineus* and *C. sinensis* and second to compare the complements for similarities and differences in expression and most importantly in sequential composition. Finally we describe expression differences in *O. felineus* stages for both miRNAs and mRNAs, identifying miRNA:mRNA interactions of possible importance for the development of *O. felineus.*

**Figure 1.**
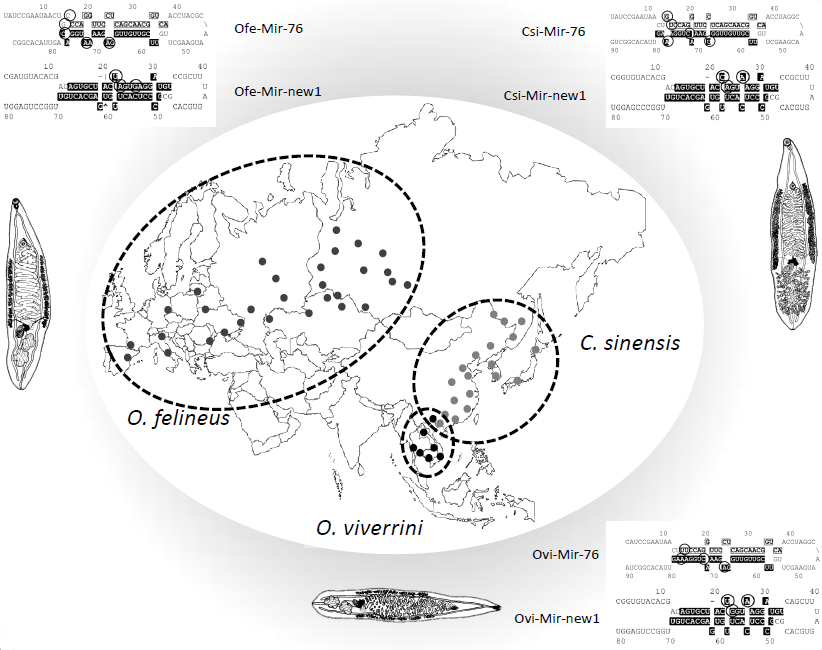
Distribution of opisthorchiids in Eurasia and how they differ in the two variable miRNA loci. Light grey dots indicate locations where *O. felineus* was detected, dark grey dots – *C. sinensis,* black dots – *O. viverrini.* Mature miRNAs are highlighted by black color. Differences in mature and star sequences are indicated by black circles.

Altogether, in total, 1 billion small RNAseq reads of sixteen published miRNA NGS datasets for all 3 opisthorchiids [11, 12], the references genomes for *O. viverrini* [14], *C. sinensis* [15] and the draft assembly of *O. felineus* (Ershov et al. in prep) were used. For RNAseq data we used annotations and expression levels directly from Pomanznoy et al [5]. Briefly, small RNAseq reads of the projects PRJNA270708 [11] and PRJNA127731 [12] were downloaded from the Sequence Read Archive (SRA) and processed as described before [11, 16]. Genomic references were downloaded from http://parasite.wormbase.org/ (PRJDA72781 & PRJNA222628) and made available by Ershov et al respectively. Using the miRNA prediction algorithm MirMiner [17] & (Fromm et al in prep) and applying a consistent set of criteria for the annotation of miRNA genes [18], we reanalyzed and reannotated opisthorchiids’ miRNAs.

We found that the miRNA complements of the three opisthorchiids are very similar and much larger than presumed: they are composed of 55 conserved miRNAs (34 families) shared by the three flatworms and only found support for 1 novel miRNA (Supplementary tables 1 and 2, Supplementary file 1). Our prediction algorithms found 35 previously missing conserved miRNA genes that belong to 22 conserved miRNA families and 1 novel miRNA gene. Further we rejected 96 previously described novel miRNA genes because they did not fulfill annotation criteria for *bona fide* miRNA genes [18] (Supplementary table 3). A noteworthy finding is that Mir-76 and the Mir-Novel-1 show sequential differences between the 3 species while having abundant expression levels (Figure 1).

When we compared the expression patterns of all miRNAs in the adult worm datasets of all three species we found that they are very similar, too. The top three expressed miRNAs in adults of all three species were Mir-10-P2a, Mir-71-P1 and Mir-281. It is worth noting that we were not able to detect expression of Mir-12 in *O. viverrini* and *O. felineus* but because the sequence is identical to the very little expressed version of Mir-12 in *C. sinensis* we included it for both species, too (Supplementary table 2, asterisks).

Nevertheless, the homogenous pattern of miRNA expression we observed among the different species was not found when we compared miRNA abundance in the different stages available for *O. felineus.* We found highly distinct miRNA expression patterns between the datasets for metacercariae and the adults (Figure 2, Supplementary table 4, Supplementary figure 1). Remarkably, we were unable to detect Mir-76, Mir-10-P3 and Mir-2160-P1 in the metacercariae datasets. Because miRNA regulate gene-expression on the mRNA level we asked if we can observe a connection between the reported mRNA level differences between adult and metacercarians and the miRNA level differences we observe between the adult *O. felineus* and metacercarian stages. Previously, the transcriptome analysis of two *O. felineus* stages identified 12,665 distinct transcripts of those 903 were metacercariae specific and 648 adult specific. In total, seven pathways were significantly enriched for differentially expressed genes (Lysosome, Neuroactive ligand–receptor interaction, Phagosome, Riboflavin metabolism, ECM–receptor interaction, Tyrosine metabolism and Arginine and proline metabolism). Consequently, we performed bioinformatics miRNA target prediction on the 3’UTR sequences of mRNA downloaded from GenBank. To ensure that we identify highly likely targets we used the intersection of three widely used programs (RNAhybrid, PITA and TargetScan) and identified 291 mRNA-targets for 46 miRNAs of *O. felineus* (Supplementary table 5).

**Figure 2.**
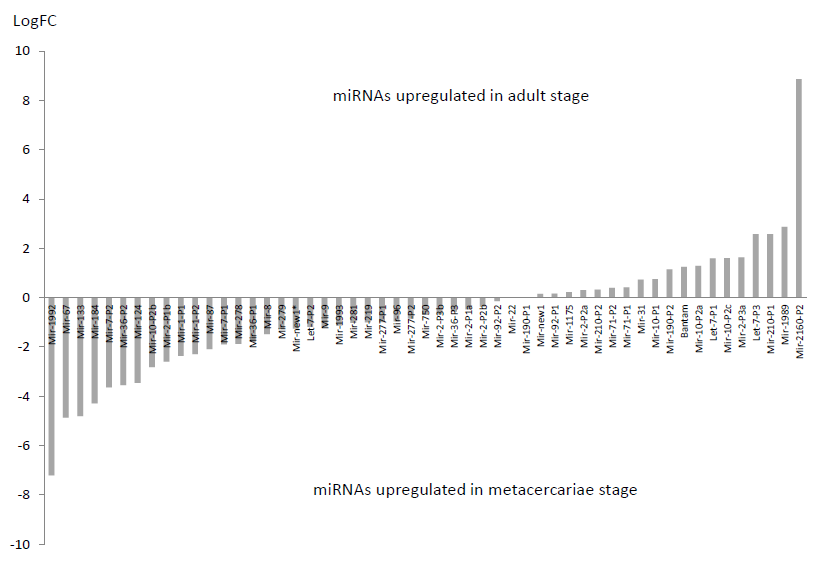
Differential expression of miRNAs between adult and metacercariae stages.

We analyzed the predicted targets of adult specific miRNAs (Mir-76, Mir-10-P3 and Mir-2160-P1), five most upregulated (Mir-2160-P2, Mir-1989, Mir-210-P1, Let-7-P3 and Mir-2-P3a) and five most downregulated miRNAs (Mir-1992, Mir-67, Mir-133, Mir-184 and Mir-7-P2) in the adult stage. All targets that did not follow prediction were excluded from further analysis. Of the 61 predicted targets for the 13 enriched miRNAs (3 adult specific, 5 upregulated in adult stage and 5 upregulated in metacercariae stage), 32 target mRNAs were found to behave according to a model of miRNA:mRNA interaction (Supplementary figure 2). Based on the available annotation of *O. felineus* mRNA we see several miRNAs that target metabolic processes such as transcription, DNA replication and autophagy upregulated in metacercariae and their targets are downregulated (Supplementary table 6). This is consistent with the resting state of metacercariae. Among all the downregulated targets of upregulated miRNAs in the adult stage we would like to particular mention one target of the highly deregulated Mir-2160-P2: GBJA01010536. It is a 5-hydroxytryptamine receptor 7 homolog (5HT7) and has been shown to significantly decrease motility when downregulated in *Schistosoma mansoni* adults and larvae [19]. The fact that the 5HT7 has a tenfold decreased expression in the adult *O. felineus* compared to the metacercariae requires further research. Although miRNA and mRNA datasets derive from different studies, it seems that they can be used to arrive at interesting hypotheses that present the basis for further studies of the biology of trematode development.

Altogether, we find that putative variation of microRNAs was an artifact and based on the incorrect annotation of miRNAs. Indeed, the Eurasian wide distributed opisthorchiids show extreme conservation of their miRNA complements which implies a very recent evolutionary split or even conspecificity as previously shown for monogenean parasites [20]. Further sampling of more strains is warranted in order to investigate their relationship and taxonomic status. Regardless of status, the three organisms vary in two mature miRNAs that could be used to differentiate them. The numerous newly identified conserved miRNAs and their stage specific expression profiles represent potent targets for further downstream analyses, biomarker discovery and disease control.

## Acknowledgments

This work was supported by the state project of the Institute of Cytology and Genetics, the Siberian Branch of the Russian Academy of Sciences, [project No. 0324-2016-0002]. B.F. was supported by the South-Eastern Norway Regional Health Authority grant #2014041. The authors thank Michael Hackenberg for help with data acquisition from the SRA. The Siberian Branch of the Russian Academy of Sciences (SB RAS) Siberian Supercomputer Center is gratefully acknowledged for providing supercomputer facilities.

